# Preclinical assay of the effects of lacosamide, pregabalin and tapentadol on the rat N1 spinal somatosensory evoked potential

**DOI:** 10.1101/2025.04.25.650582

**Authors:** Kenneth A J Steel, James P Dunham, Caterina Leone, Andrea Truini, Rolf-Detlef Treede, Keith Geoffrey Phillips, Jeffrey Krajewski, Anthony Blockeel, Anthony E. Pickering

## Abstract

The high failure rate in translating novel analgesics into the clinic has highlighted the need for more translatable biomarkers of analgesic efficacy. The N13 component of spinal somatosensory evoked potential (SEP) has been proposed as a biomarker of spinal nociceptive processing in humans, but it is not known whether this can be back translated into rodents. Tapentadol, lacosamide and pregabalin were used as pharmacological probes to assess the sensitivity of spinal SEPs to drug action.

In anaesthetised, naïve rats (n=44), a multielectrode silicon probe was inserted into the L4 spinal cord to record SEPs from the dorsal horn following electrical stimulation of the sciatic nerve. At baseline, the N1 component (rodent equivalent of the human N13) had an amplitude of 1.33 ± 0.07mV at a latency of 4.6 ± 0.2ms following low-intensity stimulation, with an intensity-dependent amplitude increase into the noxious range.

The N1 amplitude was significantly reduced by 10mg/Kg tapentadol (40.2 ± 12.5 % vs vehicle 96.2 ± 8.0 %) and 30mg/Kg lacosamide (46.3 ± 20.9 % lacosamide vs vehicle 115 ± 5.9 %) at 60 minutes after intraperitoneal administration. Tapentadol also reduced the N1 amplitude in the noxious range. Lacosamide increased the stimulus current required to evoke the half maximal N1 response (EC50), without reducing the maximum N1 amplitude in the noxious range. Pregabalin (at any dose up to 30mg/kg) did not modulate the N1 amplitude.

These results show the spinal N1 is differentially modulated in a way that reflects distinct mechanisms of drug action consistent with it being a translatable biomarker of analgesic efficacy.

## 1. Introduction

An important factor in the limited success in transitioning novel analgesics into clinical use is the mismatch between apparent efficacy in animal pain models and the successful treatment of human pain conditions [35,39]. Behavioural observations and neural recordings from animal models serve as primary tools for evaluating analgesic efficacy before human trials [38], despite their limited face validity in replicating clinical conditions. It has been suggested that introducing functional biomarkers of target engagement into research programs could facilitate future clinical success [18,51,62,69]. Such biomarkers should be capable of assaying conserved pathways within the nociceptive system, strengthening findings from preclinical experiments.

One potential biomarker of nociceptive processing in the spinal cord is the N13 component of the somatosensory evoked potential (SEP) [45]. Recorded above the sixth cervical vertebra, the N13 represent neuronal activity from the spinal dorsal horn mediated by Aβ-fibres following electrical stimulation of the medial nerve [5,16,30,45,46,53]. Human studies have shown the amplitude of the N13 is differentially modulated in a variety of experimental pain conditions [42]. Topical capsaicin was shown to increase the amplitude of the N13, and this effect was attenuated by pregabalin [44], a drug used in chronic neuropathic pain management [19,64]. The amplitude of the N13 is also reduced by conditioned pain modulation [54], and it has been suggested that this is mediated through endogenous modulation of wide dynamic range (WDR) neurons [2–4,41,54]. WDR neurons respond to both innocuous and noxious stimuli, as they receive convergent inputs from Aβ-fibres, Aδ-fibres and C-fibre nociceptors [50,57], and provide a rationale for why the innocuously evoked N13 could be used to monitor changes in nociceptive processes. A rat equivalent of the N13 (named the N1 component) can be recorded under anaesthetic from with dorsal horn [1,28,52,56]. However, it remains unclear whether the amplitude of the rodent N1 is modulated by analgesics.

This study will investigate how the rodent N1 is modulated by *standard-of-care* analgesics to assess its potential as a functional biomarker of the nociceptive system and its modulation. The drugs were selected because of their similar pharmacokinetic characteristics (short time to peak-concentration, lack of metabolites and relatively short half-life) and the fact they act on different compartments of the nociceptive system [51]. Lacosamide, typically an anti-epilepsy treatment, has an inhibitory action on peripheral voltage gated sodium channels [7]. Pregabalin, an anticonvulsant, targets the α_2_δ-subunit of N-type calcium channels at a spinal level [19]. Tapentadol, an analgesic with an opioidergic and noradrenergic reuptake inhibition dual mechanism of action in the brain and spinal cord [6]. In anaesthetised rats, a multi-electrode was inserted into the lumbar spinal dorsal horn to record SEPs following electrical stimulation of the sciatic nerve at graded intensities from the innocuous to the noxious range. The activation of different primary afferent classes, dependent on their activation thresholds, produced additional SEP components corresponding to their conduction latencies. Ultimately, the results of this study will be compared against concurrent human data collected by collaborators within the IMI Pain Care Consortium [43].

## 2. Methods

Experiments were performed in accordance with the UK Animals (Scientific Procedures) Act 1986. All experiments were approved by the University of Bristol Animal Welfare and Ethical Review Board. Male Wistar rats (n=44, Envigo, UK) weighing between 225g-375g were housed in cages under a 12-hour alternating light/dark cycle, and with *ad libitum* access to food and water.

### 2.1 Drugs

Drugs were prepared fresh on the day of each experiment. Each rat received an intraperitoneal injection, volume 10ml/kg, of either lacosamide (vehicle, 3 mg/kg, 10 mg/kg & 30 mg/kg), pregabalin (vehicle, 3 mg/kg, 10 mg/kg & 30 mg/kg), or tapentadol (vehicle, 3 mg/kg &10 mg/kg). Tapentadol was not dosed at 30mg/kg as it caused deterioration in the physiological condition of anaesthetised rats due to respiratory depression. Vehicle solutions were prepared as stocks and in each case the drug was dissolved fresh from powder (lacosamide and pregabalin were supplied by Novachemistry, UK and tapentadol was supplied by Precise Chemipharma, India):

- Lacosamide - in a 1% hydroxyethyl cellulose, 0.5% TWEEN®80, 0.05% Antifoam SE-15 (1% HEC) solution. This carrier was formulated by dissolving HEC into 250ml dH_2_O at 70^°^C before adding TWEEN®80 and Antifoam SE-15 and stirring for an additional 30 minutes at 70°C [8].
- Pregabalin - in 0.5 % hydroxypropyl methylcellulose (0.5% HPMC) solution which was initially dissolved by stirring in dH_2_O at 70^°^C for 30 minutes [55].
- Tapentadol, in 0.9% saline.

### 2.2 Randomisation and blinding

Experiments were performed in a block-randomised, blinded manner, such that animals were first allocated to treatments (first block) and then dose (second block; Table 1). On the day of the experiment, a colleague not involved with the study, selected a vial of drug solution to be administered. The experimenter was blinded to the dose, but not to the drug, because of the different recording times needed for the drugs (60 minutes for Tapentadol and Lacosamide, and 90 minutes for pregabalin due to a longer time to maximal concentration (Tmax) [26,59,67]). Blinding was maintained until data analysis was completed. Each animal received a single dose of drug and group sizes were n=4 rats for each dose of drug.

**Table 1.**
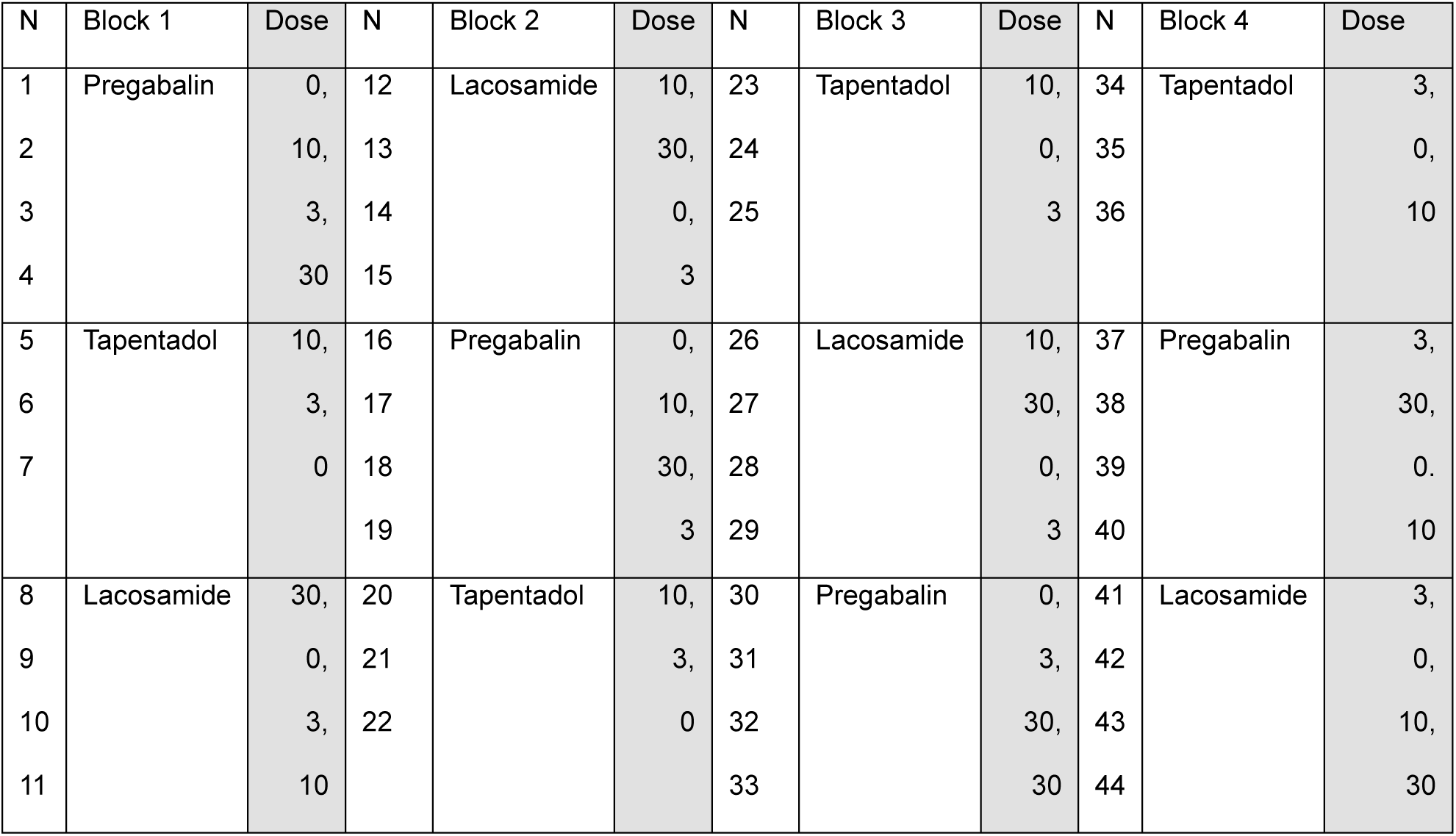
Example of block-randomisation. Experiments were organised into blocks containing each dose of drug. The order of drugs and dosing were randomly assigned within blocks. On each day of the experiment, the animal (N) was dosed with the corresponding drug and dose.

| N | Block 1 | Dose | N | Block 2 | Dose | N | Block 3 | Dose | N | Block 4 | Dose |
| --- | --- | --- | --- | --- | --- | --- | --- | --- | --- | --- | --- |
| 1 | Pregabalin | 0, | 12 | Lacosamide | 10, | 23 | Tapentadol | 10, | 34 | Tapentadol | 3, |
| 2 |  | 10, | 13 |  | 30, | 24 |  | 0, | 35 |  | 0, |
| 3 |  | 3, | 14 |  | 0, | 25 |  | 3 | 36 |  | 10 |
| 4 |  | 30 | 15 |  | 3 |  |  |  |  |  |  |
| 5 | Tapentadol | 10, | 16 | Pregabalin | 0, | 26 | Lacosamide | 10, | 37 | Pregabalin | 3, |
| 6 |  | 3, | 17 |  | 10, | 27 |  | 30, | 38 |  | 30, |
| 7 |  | 0 | 18 |  | 30, | 28 |  | 0, | 39 |  | 0. |
|  |  |  | 19 |  | 3 | 29 |  | 3 | 40 |  | 10 |
| 8 | Lacosamide | 30, | 20 | Tapentadol | 10, | 30 | Pregabalin | 0, | 41 | Lacosamide | 3, |
| 9 |  | 0, | 21 |  | 3, | 31 |  | 3, | 42 |  | 0, |
| 10 |  | 3, | 22 |  | 0 | 32 |  | 30, | 43 |  | 10, |
| 11 |  | 10 |  |  |  | 33 |  | 30 | 44 |  | 30 |

### 2.3 Surgery and preparations for spinal electrophysiological recording

Rats weighing 225-375g were anaesthetised with isoflurane until loss of paw withdrawal reflex (5% isoflurane at 1.0l/min for induction and maintenance at 2% at 0.5l/min in O_2_). A 25-gauge needle attached via small-diameter tubing using standard Luer connectors was bent and carefully inserted into the intraperitoneal cavity for drug administration [14]. The skin over the thoracolumbar spine was shaved and sterilised with chlorohexidine wipes. The rat was transferred into the stereotaxic frame (David Kopf Instruments, Canada) and core body temperature of 37°C was maintained using a heat blanket with rectal thermometer attached to a homeothermic monitor (Harvard Apparatus, USA).

After a skin incision at T13-L3, the connective tissue and paraspinal muscles were removed to expose the spinous processes. A laminectomy was performed to expose the L4-L5 segments of the spinal cord. The spine was then stabilised using vertebral clamps on the intact superior and inferior spinous processes and notch clamps perpendicular to the delaminated vertebra (Figure 1A-). A small section of spinal dura was removed with a hooked needle to create a hole for the probe and mineral oil was applied to prevent the tissues from drying.

**Figure 1.**
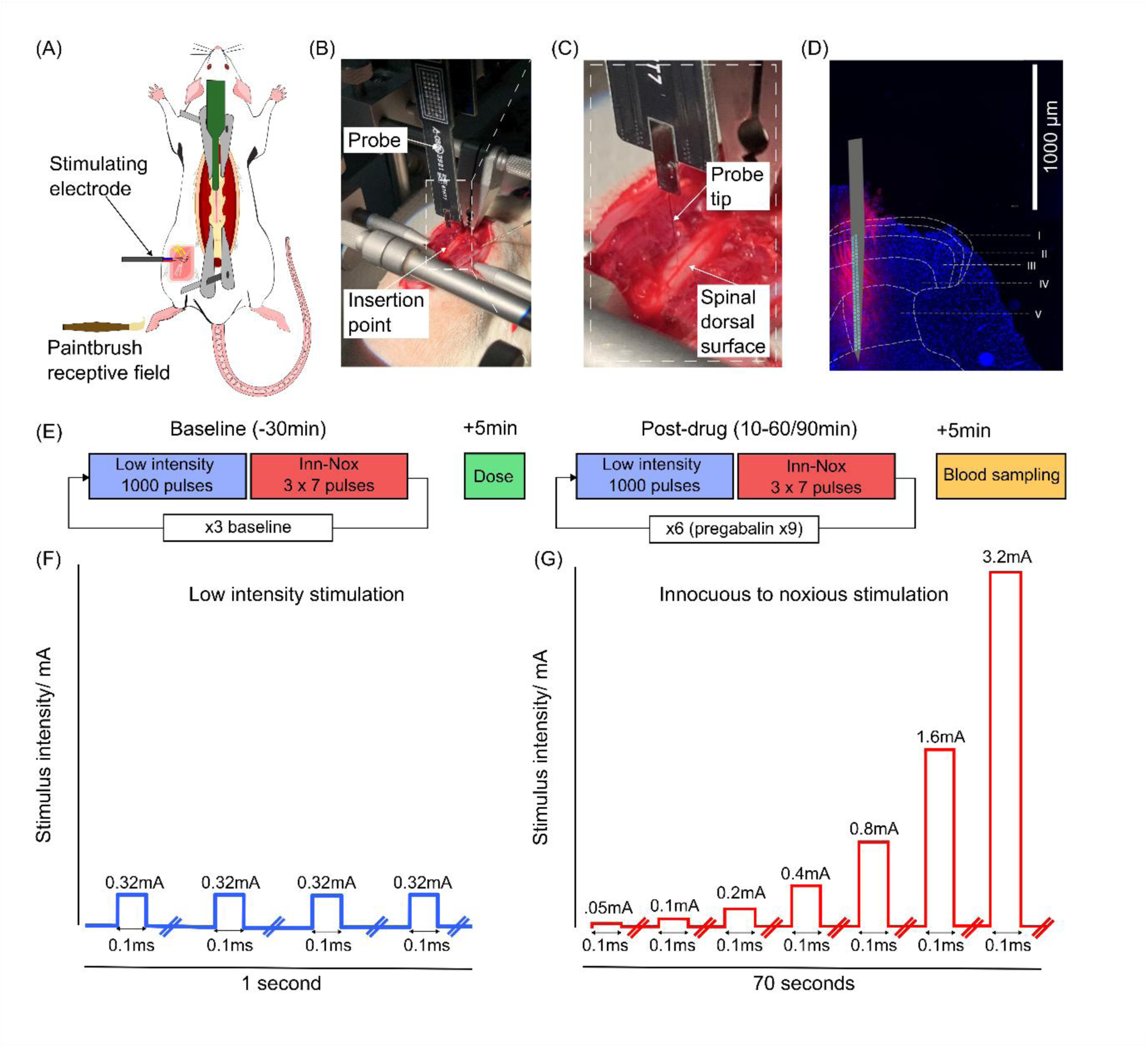
Overview of experimental approach. (A) Recordings were made from the lumbar (∼L4/5) spinal dorsal horn in anaesthetised male Wistar rats implanted with stimulation electrodes on the sciatic nerve (B) A laminectomy of T13-L1 vertebra exposes the dorsal surface of the spinal cord, providing access for the recording electrode. (C) Close-up of probe insertion - the dura mater is opened and covered in mineral oil before the probe is carefully inserted into the tissue. (D) Coronal lumbar spinal cord section stained with DAPI and DiI lipophilic tracer marking the placement of the probe within the dorsal horn. A schematic of the multi-electrode probe has been superimposed. (E) The SEP assay protocol; 3 repeats of the stimulation paradigm were used to characterise spinal SEPs at baseline. 5 minutes post dose, the stimulus paradigm was resumed and continuously repeated until the end point. Terminal blood samples were taken 5 minutes after the end point. (F) Low intensity stimulation paradigm; 1000 repeats of rectangular electrical pulses with a constant current (= 1.5x paw twitch threshold, average 0.32mA) and 0.1ms pulse width. 4Hz frequency, // = 250ms. (G) Inn-Nox stimulation paradigm; 3x7 repeats of rectangular electrical pulses starting a 0.05mA and doubling in current until 3.2mA. 0.1Hz frequency, // = 10s.

A small incision was made above the left bicep femoris muscle, and the sciatic nerve was exposed using careful blunt dissection proximal to its trifurcation. A tissue retractor was used to construct a pool for mineral oil to prevent the nerve from drying and to improve access for the stimulator electrode. A custom bipolar Ag-AgCl hook electrode (Dr. G. Newton) connected to a constant current stimulator (DS4, Digitimer, UK) was used to electrically stimulate the sciatic nerve (Figure 1A). The interval and intensity of the single-square current pulses were controlled from Spike 2 (Cambridge Electronic Design, UK) and were recorded in Open Ephys.

To record neural activity, a 64-channel silicon probe (Cambridge Neurotech ASSY-77 H5) was used in combination with 2x 32 channel head stage amplifiers (Intan, RHD2132, #0415 & #0917) connected using a Neuronexus 64 channel head stage adaptor. Signal from the probe was sampled at 30kHz with an Open Ephys acquisition board and software (V0.6.0.1). Using a motorised manipulator (Scientifica, UK), the probe was inserted slowly ∼800μm into the spinal dorsal horn and left to stabilise for 10 minutes (Figure 1C). In some experiments, the probe was dipped into DiI (1′-dioctadecyl-3,3,3′,3′-tetramethylindocarbocyanine; Thermo Fisher Scientific, UK) and left to dry for 10 minutes before insertion to stain the recording site. The data were visualised in the Open Ephys graphical user interface and bandpass filtered simultaneously at 1-300Hz and 600-6000Hz to observe both local field potentials and single unit action potentials respectively. Successful probe insertion in a segment of the dorsal horn receiving hind paw afferent inputs was confirmed by the profile of spontaneous action potentials across multiple channels along the length of the probe and evoked action potentials following light brush of the hind paw.

To best mimic the N13 potential SEPs were initially generated by electrical stimulation of the sciatic nerve (stimulus duration: 0.1ms; frequency: 0.1Hz) at 1.5x the intensity threshold for an evoked twitch in hind paw [37]. The depth of the probe was adjusted to ensure the amplitude of the N1 component was maximal in a central region of the probe so the peak amplitude of the SEP could be tracked across electrode contacts in the event of movement of the spinal cord over the course of the recording.

### 2.4 Somatosensory evoked potential assay protocol

Once the probe was positioned correctly, a standard protocol was started to characterise the SEP (Figure 1E). The protocol was designed to track the amplitude of the N1 component in response to low and high-intensity electrical stimulation. Recordings consisted of a 30-minute baseline period and a 60/90-minute post-dose period:

1. A 2-minute recording of intermittent light brushing of the receptive field.
2. A 30-mininute baseline recording of SEPs, consisting of 3 repeats of a 10 min stimulation paradigm. This 10 minute stimulation paradigm included:

- Low-intensity electrical stimulation (1000 x 0.1ms pulses at 4Hz,1.5x threshold to evoke a hind paw twitch).
- 1 min rest period.
- Ramping electrical stimulation from the innocuous to the noxious range (Inn-Nox stimulation; 3 repeats of 7 x 0.1ms pulses at 0.1Hz, from 0.05mA doubling in intensity until 3.2mA).
- 1 min rest period.
3. Administration of either vehicle or drug solution (10ml/Kg) delivered via the i.p catheter.
4. A 5 min pause.
5. Up to 60/90-minutes post-dose, continuous repeats of the 10 minute stimulation paradigm detailed in step 2.
6. Blood sampling for pharmacokinetic-pharmacodynamic modelling at 65- or 95-minutes post dose. Blood samples were collected from a small incision of the hind paw by pipetting spots onto filter paper.

### 2.5 Drug concentration quantification

Dried blood spot (DBS) analysis was performed by Q^2^ Solutions (North Carolina, USA) after all experiments were completed. DBS samples were analysed for lacosamide, pregabalin, or tapentadol concentrations using liquid chromatography-tandem mass spectrometry (LC-MS). A 3 mm punch from each DBS was placed in wells of a 96-well plate, and 180 µL of an acetonitrile-water solution (1:1) was added to facilitate analyte extraction. The plate was shaken for 45 minutes at room temperature to maximize the extraction of analytes. Following extraction, samples underwent a two-fold dilution with a formic acid and water buffer to enhance compatibility with the LC-MS system. Analytes were separated using a Shimadzu LC system with LC-30AD pumps, with an initial flow rate of 1.5 mL/min. Mass spectrometric analysis was conducted using an AB Sciex API 4000 triple quadrupole mass spectrometer (Massachusetts, USA), interfaced with the Shimadzu LC system. Standard concentration curves were generated by analysing target analytes with a known concentration, enabling sample quantification based on peak area data. Final concentrations of the drugs were calculated using the drugs relative molecular weight (tapentadol 257.8 g/mol, lacosamide 250.29 g/mol, and pregabalin 159.23 g/mol).

### 2.6 Data analysis and statistics

All data were analysed in MATLAB (R2022a) and GraphPad Prism 10. The statistical analysis plan was pre-published on the Open Science Framework and contains the custom MATLAB scripts used (https://osf.io/ghf8p/?view_only=8d000c3a26f448a8a1088d461d86c490). This is described in brief below:

Data from each electrode contact and event markers were imported to MATLAB and the data were low pass filtered at 300Hz to remove stimulus artefacts. Events were separated into 10-minute epochs, corresponding to the 10minute stimulus protocol (3 baseline + 6/9 post dose) across the duration of the experiment and grouped according to their stimulus intensity (low-intensity stimulation and innocuous-noxious stimulation). For each channel, 300ms samples of data around each electrical stimulus (−100ms to +200ms) were extracted to capture all SEPs. The sampled data were then averaged across repeated stimuli within each epoch to produce an averaged SEP waveform for the low-intensity stimulation and Inn-Nox stimulation. The amplitude of the N1 component was extracted from the first negative peak of the SEP. To track the amplitude of the N1 across the time course, the amplitude of the N1 component within each epoch was selected from the electrode site that recorded the N1 component with the peak amplitude. Secondary and tertiary negative components of the SEP (N2 and N3) were visually identified from the averaged waveforms. The conduction velocity (m/s) of the N1 component was estimated by dividing the distance between stimulation and recording site, 8cm, by the latency of the N1 component. Data from one animal were excluded from the study because the peak of the SEP moved abruptly across the electrode contacts (indicating mechanical displacement) and therefore could not be tracked.

Data are presented as mean normalised N1 values relative to the baseline (100*value / mean of first 3 time points) ± standard error of the mean (SEM). Statistical significance for the primary endpoint (percentage change from baseline of N1 amplitude) and the stimulus-response curve were calculated using mixed-effects analysis with Dunnett’s multiple comparisons test. Responses from the Inn-Nox stimulation paradigm were analysed using least squares regression model to give the EC50 (effective current required for half the maximal) of the N1. Statistical significance for differences between groups in the EC50 were calculated using the extra sum-of-squares F test. Dose-response data were calculated from the end-point amplitude of the N1 component to low-intensity stimulation vs blood plasma concentration and were analysed using least squares regression model to fit a dose-response curve.

### 2.7 Histology

In experiments where DiI was used to stain the recording site, the animals were transcardially perfused with phosphate-buffered saline (PBS) and the spinal cord was removed via hydraulic extrusion with PBS (for methods, see (Richner et al., 2017)). The spinal cord was fixed in 4% formaldehyde solution for up to 24 hours before being transferred to a 30% sucrose solution for at least 48 hours. The lumbar spinal segment was cut into 50μm sections using a freezing microtome and transferred into PBS solution before being mounted in series onto glass slides. The mounted tissue was left to dry for 24 hours, before the neuronal cell bodies were stained using 300nM DAPI (4′,6-diamidino-2-phenylindole) and the sections cover slipped.

Fluorescence microscopy was performed (Lecia DM 4000B microscope, Wetzlar Germany) under ultraviolet light (395nm) for the DAPI staining and under green illumination (564nm) for the DiI probe marker. Captured images were merged in FIJI image processing (LOCI, University of Wisconsin) and the probe position estimated by fitting a line through the apical points of the DiI fluorescence. Spinal dorsal horn laminae were marked by reference to the Atlas of the Rat Spinal Cord ([66], Figure 1D).

## 3. Results

### 3.1 Baseline characterisation of the SEP

For each animal, low-intensity electrical stimulation of the sciatic nerve was titrated to be 1.5x threshold to evoke a hind paw twitch (0.32±0.03 mA). This generated a spinal SEP that was recorded from the 64-channel multielectrode probe inserted into the spinal dorsal horn (Figure 1D). The SEP was observed in the proximal and mid-range channels of the probe (corresponding to Lamina IV-V, Figure 2A). The N1 component had an average amplitude of 1.33 ± 0.07mV and an average latency of 4.6 ± 0.2ms (n=44). The latency of the response was used to estimate the conduction velocity of 22.2m/s for the activated peripheral axons (distance of 0.08m, allowing 1ms for synaptic transmission from the primary afferent), which is consistent with the conduction velocity of rat Aβ-primary afferent neurons [32].

**Figure 2.**
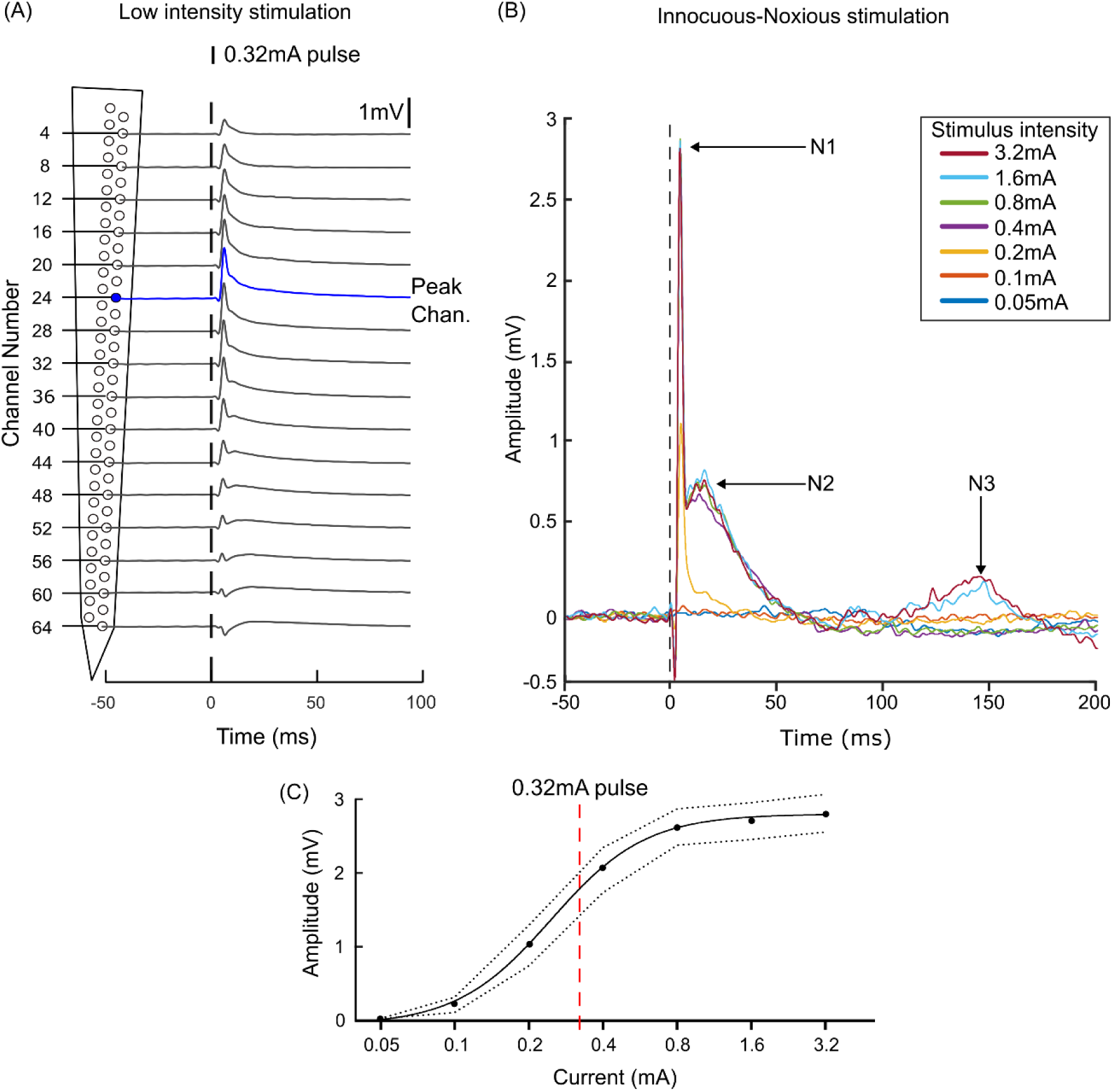
Baseline characterisation of the rat lumbar spinal somatosensory evoked potential. A. Representative averaged waveforms of the spinal SEP to low-intensity stimulation along the length of the probe. In this example, channel 24 recorded the SEP with the highest amplitude N1 component (blue). Note: the SEP is presented as inverted as is convention when reporting negatively deflecting potentials in clinical studies. B. Superimposed traces of the spinal SEP following innocuous-noxious (Inn-Nox) electrical stimulation. In this example, the N1 component was evoked at stimulus intensities ≥0.1mA (see orange trace) and its amplitude plateaued at stimulus intensities ≥0.8mA. The N2 component was evoked at stimulus intensities ≥0.2mA and plateaued at stimulus intensities ≥0.8mA. The N3 component was only evoked at 1.6mA and 3.2mA stimulus intensities. C. Stimulus response curve for the amplitude of the N1 component to Inn-Nox electrical stimulation (n=12). The mean, titrated stimulation intensity used for the low-intensity stimulation paradigm marked in red.

Following Inn-Nox stimulation, the amplitude of the N1 component was modulated in an intensity dependent manner (Figure 2B). This increase in the amplitude of the N1 component was accompanied by the appearance of additional negative components named N2 and N3, which respectively reflects the intensity-dependent recruitment of higher threshold Aδ- and C-fibre primary afferents [5,37]. The stimulus-response curve for the N1 component showed the electrical threshold for generation was 0.1mA, the EC50 of the response was 0.25mA and the peak amplitude plateaued above 0.8mA (Figure 2C).

### 3.2 Tapentadol

Tapentadol was administered to assess the effect of a mixed mu-opioid and noradrenaline re-uptake inhibitor on the N1 component. Following baseline characterisation of the SEP, tapentadol (3, 10mg/Kg and vehicle (n=4 per group)) was injected and the amplitude of the N1 component was tracked for 60 minutes. Tapentadol altered the amplitude of the N1 evoked by low-intensity stimulation (mixed-effects analysis showed a significant effect of time (P<0.0001, F= 7.792), of dose (P=0.0148, F=6.985) and a dose x time interaction (P<0.0001, F=3.868)). Post-hoc analysis showed tapentadol at 10mg/kg reduced the amplitude of the N1 component (compared to vehicle) after 10 minutes (103.5 ±4.3% vs 58.7 ±12.0%, P= 0.0042), 20 minutes (101.6 ±8.2% vs 53.4 ±24.3%, P= 0.0073), 30 minutes (101.6 ±8.1 vs 43.5 ±12.0 %, P= 0.0003), 40 minutes (102.4 ±9.4 vs 37.5 ±12.8 %, P<0.0001), 50 minutes (98.3 ±11.5 % vs 39.5 ±13.3%, P=0.0002), and 60 minutes (96.2 ±8.0 % vs 40.2 ±12.5 %, P=0.0003) post-injection (Dunnett’s multiple comparisons test; Figure 3A & 3B, Table 2).

**Figure 3.**
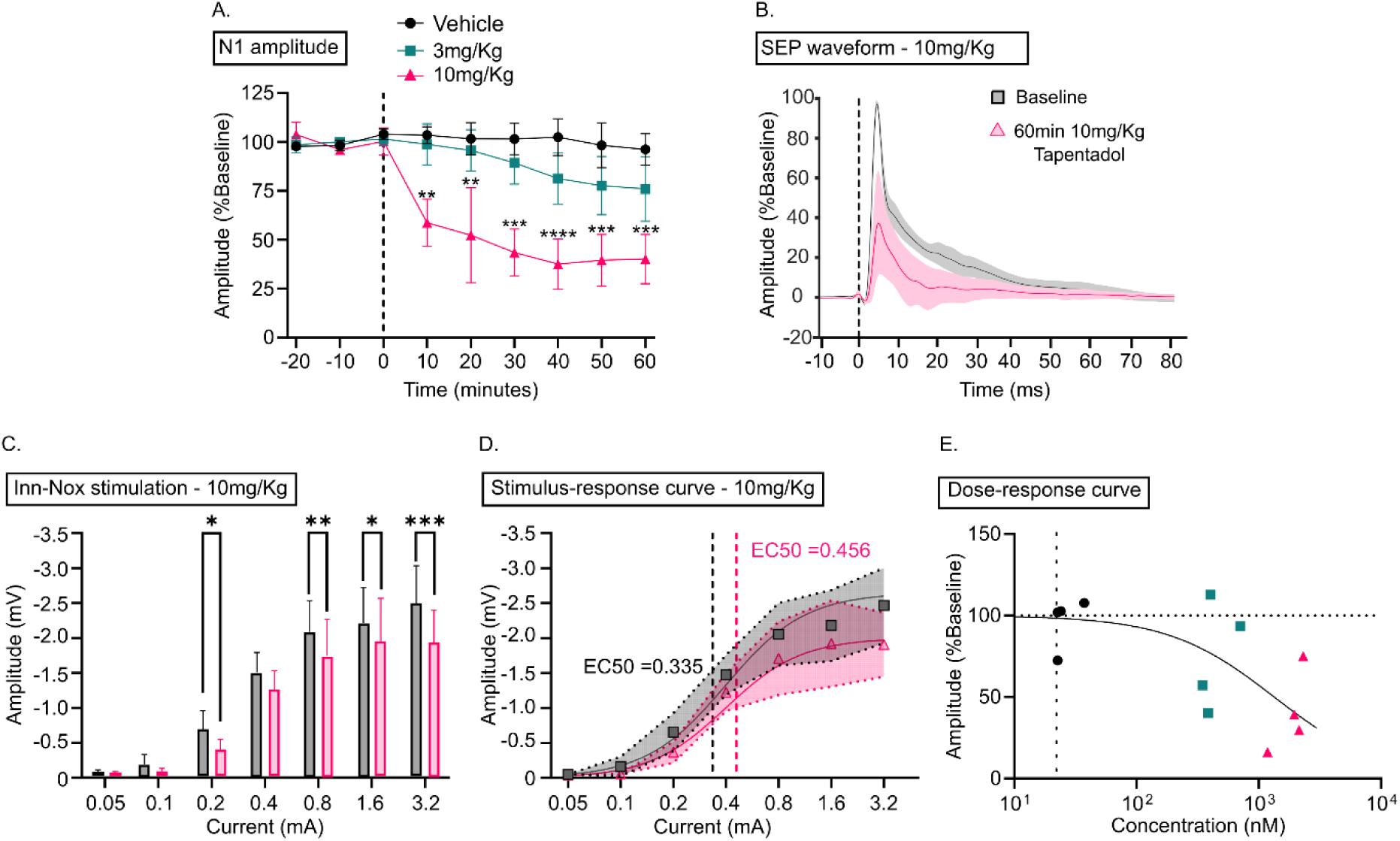
Tapentadol significantly reduces the amplitude of the N1 component to innocuous and noxious electrical stimulation, without significantly affecting the EC50 of the response. A. Effect of tapentadol dose on the baseline normalised amplitude of N1 component evoked by low-intensity electrical stimulation. Mixed effects analysis with Dunnett’s multiple comparisons test showed a significant effect of 10mg/Kg tapentadol on the amplitude of the response at each time point after dosing. B. 10mg/Kg tapentadol significantly reduced the amplitude of the SEP waveform to low-intensity stimulation at 60-minutes (pink, 40.2 ±12.6%) when compared to baseline (grey). C. Amplitude of N1 component at baseline and 60-minutes following Inn-Nox stimulation. 10mg/Kg tapentadol significantly reduced the amplitude of the N1 SEP at 0.2mA, 0.8mA, 1.6mA and 3.2mA stimulus intensities. D. Stimulus-response curves for the amplitude of the N1 component following innocuous-noxious (Inn-Nox) stimulation, with lines of best fit for baseline (grey) and 60 minutes (pink) responses (Hillslope equation). Effective current 50% (EC50) for the dose response curves for plotted as intersecting lines. E. Dose response curve of tapentadol plasma concentration at 60 minutes vs N1 amplitude evoked by low-intensity electrical stimulation, dotted line indicates sensitivity limit of the assay <22.6nM. Data are presented as the mean and normalised mean ±SEM, *P<0.05, **P<0.005, ***P<0.0005, ****P<0.0001.

**Table 2.** Tapentadol, lacosamide and pregabalin induced modulation of the amplitude of the N1 component. Data are expressed as mean amplitude (mV) ±SEM.

| Drug | Timepoint | N1 amplitude (mV) |  |  |  |
| --- | --- | --- | --- | --- | --- |
|  |  | Vehicle | 3mg/Kg | 10mg/Kg | 30mg/Kg |
| Tapentadol | Baseline | 0.93±0.2 | 1.36±0.2 | 1.05±0.2 | - |
| | 60min | 0.88±0.3 | 1.24±0.6 | 0.48±0.2 | - |
| Lacosamide | Baseline | 0.59±0.1 | 1.73±0.4 | 1.51±0.2 | 1.51±0.3 |
| | 60min | 0.70±0.2 | 1.43±0.3 | 1.57±0.5 | 1.00±0.6 |
| Pregabalin | Baseline | 1.25±0.2 | 1.12±0.2 | 1.58±0.1 | 1.57±0.2 |
| | 60min | 1.19±0.4 | 1.03±0.3 | 1.27±0.4 | 1.53±0.2 |

The stimulus-response relationships for the amplitude of the N1 component for the Inn-Nox protocol at baseline and 60 minutes post-dose were explored. Mixed-effects analysis in the 10mg/Kg group revealed a significant effect of stimulus intensity (P<0.0001, F= 13.77), a significant interaction between time x stimulus intensity (P=0.0167, F= 3.707), but no effect of time (P=0.0531, F= 9.64). Tapentadol significantly reduced the amplitude of the N1 component to both innocuous and noxious stimulus intensities (Tukey’s multiple comparisons test), at 0.2mA (0.66 ±0.27mV vs 0.37 ±0.15mV, P=0.038), 0.8mA (2.1 ±0.45mV vs 1.71 ±0.53mV, p=0.006), 1.6mA (2.18 ±0.51mV vs 1.92 ±0.61mV, p=0.0112) and 3.2mA (2.470 ±0.54mV vs 1.91 ±0.46 mV, p=0.00104; Figure 3D). Stimulus response curves were fitted to estimate the EC50 which was not significantly altered by tapentadol 10mg/kg (0.335 mA at baseline vs 0.459 mA at 60 minutes, P= 0.4, F= 0.71, Extra sum-of-squares F test; Figure 3D).

The dose-response curve of the tapentadol plasma concentration vs the amplitude of the low intensity evoked N1 component at 60min showed an effect with plasma concentrations that approached the micromolar range, with a mean concentration of 1.9 ±0.25µM in the 10mg/Kg group (Figure 3E).

### 3.3 Lacosamide

Lacosamide was used to assess the sensitivity of the N1 component to an anti-neuropathic pain medication with a mechanism of action on sodium channels (n=15; 4 x 3mg/Kg, 4 x 10mg/Kg & 4 x 30mg/Kg, 3 x vehicle). Mixed-effects analysis showed a significant effect of time (P=0.0353, F=2.196) and an interaction between time and dose on the amplitude of the N1 component (P=0.0015, F=2.427) but did not show an independent effect of dose (P=0.177, F=1.968) on the low intensity N1 SEP. Lacosamide (30mg/kg) reduced the amplitude of the low intensity evoked N1 component compared to vehicle at 40 minutes (111±4.5 % vs 62.6±14.6 %, p =0.0159), 50 minutes (113±6.8 % vs 55.6±16.3 %, p=0.004) and 60-minutes (115±5.9 % vs 46.3±20.9 %, p=0.0014) post injection (Figure 4A & 4B, Table 2, Dunnett’s multiple comparisons test).

**Figure 4.**
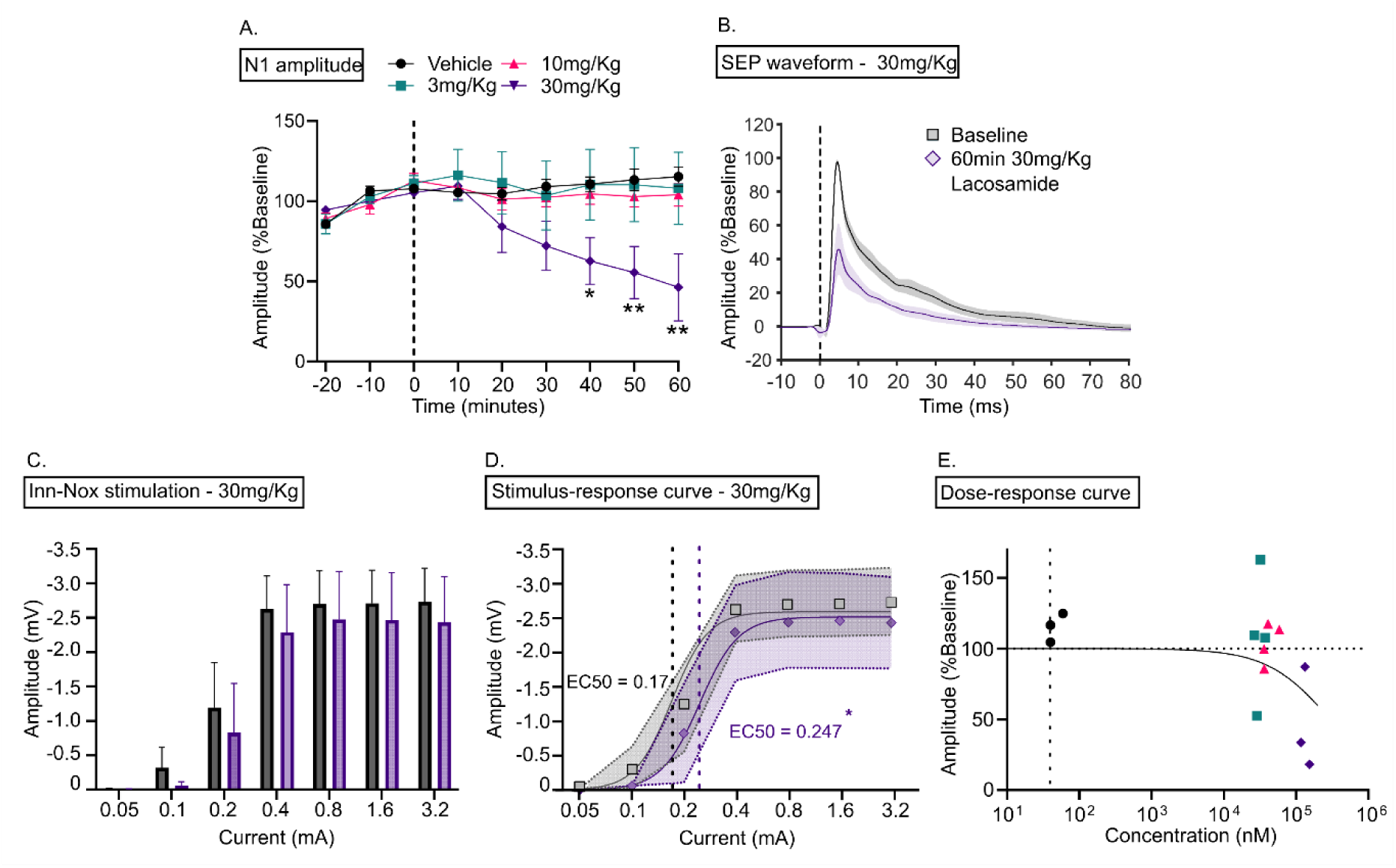
Lacosamide reduces the amplitude of the N1 component to innocuous stimulation and increases the EC50, without affecting responses to noxious stimulation. A. Effect of lacosamide on the baseline normalised amplitude of the N1 component evoked by low-intensity electrical stimulation. Mixed effects analysis with Dunnett’s multiple comparisons test showed a significant effect of 30mg/Kg lacosamide on the amplitude of the response over 40 minutes post-dose. B. 30mg/Kg lacosamide significantly reduced the amplitude of the SEP waveform to low-intensity stimulation at 60-minutes (purple, 45.61±20.2%) when compared to baseline (grey). C. The amplitude of N1 component at baseline and 60-minutes following Inn-Nox stimulation. 30mg/Kg lacosamide showed no significant effect on the amplitude of the N1 SEP to Inn-Nox stimulation. D. Stimulus-response curves for the amplitude of the N1 component following innocuous-noxious (Inn-Nox) stimulation, with lines of best fit for baseline (grey) and 60 minutes (purple) responses (Hill slope equation). The effective current 50% (EC50) was of the curve was significantly modulated by 30mg/Kg lacosamide, resulting in a rightward shift in the curve. E. Dose-response curve of lacosamide plasma concentration at 60-minutes vs N1 amplitude evoked by low-intensity electrical stimulation, dotted line indicates sensitivity limit of the assay <40nM. Data are presented as the mean and normalised mean ±SEM, *P<0.05, **P<0.005.

A two-way ANOVA of the amplitude of the N1 component following Inn-Nox stimulation at 60 minutes revealed a significant effect of stimulus intensity on the peak amplitude (P=0.0012), but no effect of lacosamide (P= 0.0529) or an intensity-time interaction (P=0.9904) when compared to baseline (Figure 4C & 4D). However, there was a rightward shift in the stimulus-response curve that was accompanied by a significant increase in the EC50 in the 30mg/Kg group at 60-minutes compared to baseline (Figure 4D, baseline 0.17 mA vs 60min 0.25 mA, P=0.0044, Extra sum-of-squares F test).

A dose-response curve of the lacosamide plasma concentration vs the amplitude of the N1 component at 60min showed the response was only modulated at plasma concentrations >100μM, with a mean concentration of 134.7 ±10.7μM (Table 3 & Figure 4E).

**Table 3.** Tapentadol, lacosamide and pregabalin terminal blood plasma concentrations. Tapentadol and lacosamide endpoint = 60 minutes, pregabalin end point = 90 minutes. Data presented as mean ±SEM.

| | Terminal blood plasma concentration (μM) | | | |
| --- | --- | --- | --- | --- |
|  | Vehicle | 3mg/Kg | 10mg/Kg | 30mg/Kg |
| Tapentadol | 0.03±0.00 | 0.46±0.08 | 1.9 ±0.25 | N/A |
| Lacosamide | 0.46±0.08 | 31.2±2.4 | 43.1±5.4 | 134.7±10.7 |
| Pregabalin | 0.06±0.00 | 15.9±4.3 | 36.5±12.0 | 145.8±10.9 |

### 3.4 Pregabalin

Pregabalin was used to assess whether the N1 component was sensitive to an anti-neuropathic pain medication with a mechanism of action on calcium channels (n=4 per group; vehicle, 3mg/Kg,10mg/Kg, 30mg/Kg). Mixed effects analysis did not reveal a significant effect of time (P= 0.125, F= 2.00), of dose (P =0.896, F= 0.197) or a time x dose interaction (P= 0.986, F= 0.518) on amplitude of the low intensity evoked N1 component (Figure 5A & 5B, Table 2).

**Figure 5.**
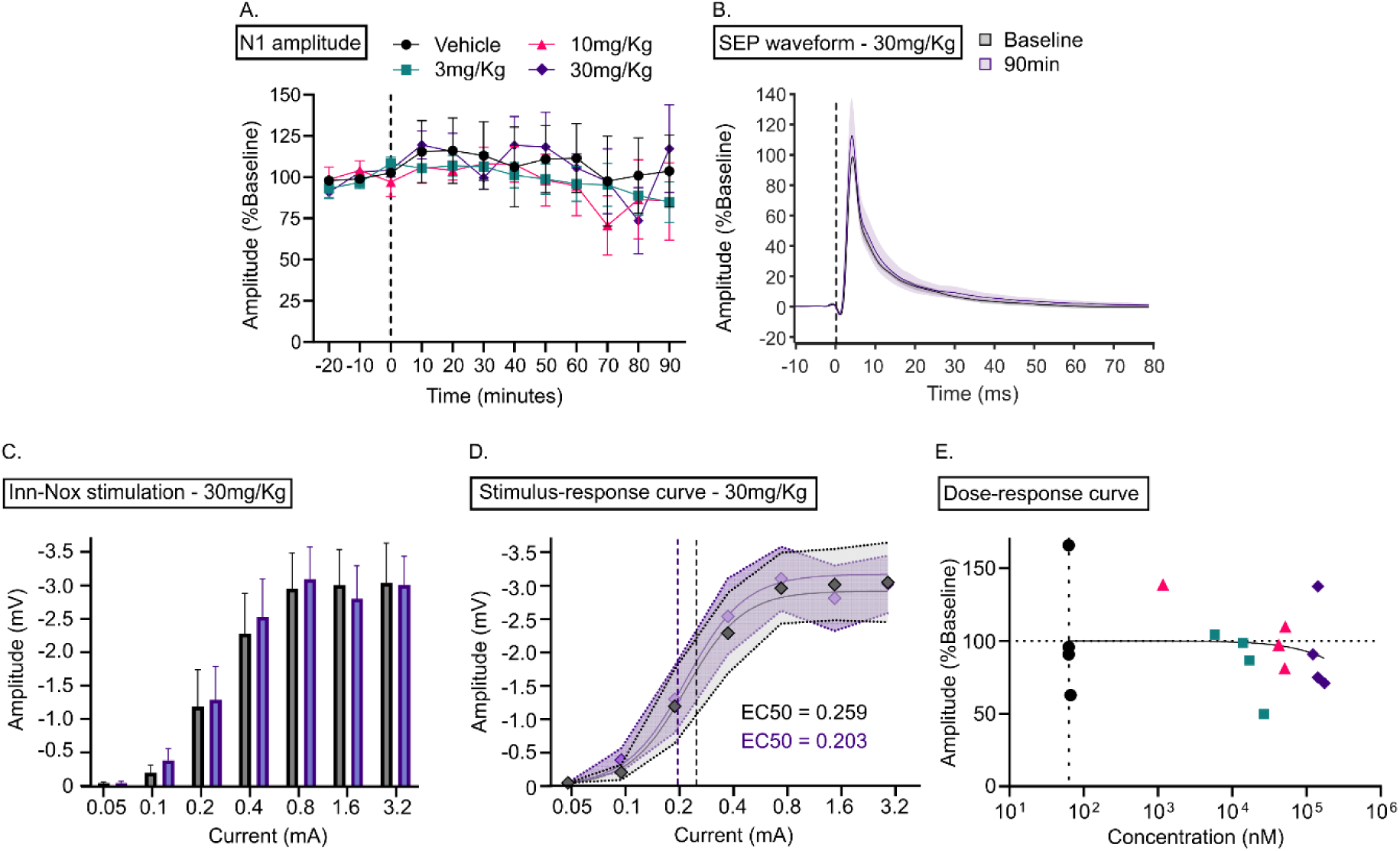
Pregabalin does not significantly modulate the amplitude of the N1 component at any concentration. A. Effect of pregabalin on the baseline normalised amplitude of the N1 component evoked by low-intensity electrical stimulation. Mixed effects analysis with Dunnett’s multiple comparisons test showed no significant effect of 30mg/Kg pregabalin on the amplitude of the response. B. 30mg/Kg pregabalin did not modulate the amplitude of the SEP waveform to low-intensity stimulation at 90-minutes (purple, 117.2 ±26.5%) when compared to baseline (grey). C. The amplitude of N1 component at baseline and 60-minutes following Inn-Nox stimulation. 30mg/Kg pregabalin showed no significant effect on the amplitude of the N1 SEP to Inn-Nox stimulation. D. Stimulus-response curves for the amplitude of the N1 component following innocuous-noxious (Inn-Nox) stimulation, with lines of best fit for baseline (grey) and 90 minutes (purple) responses (Hill slope equation). The effective current 50% (EC50) of the curve was not significantly modulated by 30mg/Kg pregabalin. E. Dose-response curve of pregabalin plasma concentration at 60-minutes vs N1 amplitude evoked by low-intensity electrical stimulation, dotted line indicates the sensitivity limit of the assay >62.9nM. Data are presented as the mean and normalised mean ±SEM.

Following the Inn-Nox stimulation, mixed effects analysis revealed a significant effect of stimulus intensity (P<0.0001, F= 19.09), but not of dose of pregabalin (P=0.503, F=0.540) or an intensity x dose interaction (P=0.0662, F=0.686), on the amplitude of the N1 component (Figure 5C & D). The EC50 of the N1 component following Inn-Nox stimulation was not significantly modulated by 30mg/Kg pregabalin at 90-minutes (baseline 0.26mV vs 0.20mV at 60min, P =0.741, Extra sum-of-squares F test; Figure 5D).

A dose-response curve generated from pregabalin plasma concentrations showed the N1 component was not modulated by the drug at any concentration up into the range of 100µM (Figure 5E & Table 3).

## Discussion

This preclinical electrophysiological study has shown that in healthy, anaesthetised rats, standard-of-care painkillers reduce the amplitude of the N1 spinal SEP. Both tapentadol and lacosamide reduced the amplitude of the N1 component evoked by low-intensity electrical stimulation and showed differential modulation of the N1 component evoked by Inn-Nox stimulation. Tapentadol reduced the N1 amplitude across the Inn-Nox stimulation paradigm, and lacosamide caused an increase in the EC50 of the response without affecting the peak amplitude. In contrast, pregabalin did not affect the amplitude of the N1 SEP evoked by any stimulus in this assay. These results show the N1 SEP is differentially modulated by these agents in a way that reflects their distinct mechanisms of action and suggest that the N1 component may represent a viable preclinical model for drug development.

The amplitude of the N1 SEP was used to assess the evoked activation of peripheral and spinal neurons as a proxy for analgesic interaction with the nociceptive system. The N1 component is thought to be generated by the simultaneous activation of spinal interneurons following the stimulation of cutaneous Aβ-primary afferents [5,30]. Using multi-electrode silicon probes, this study has replicated previous recordings of spinal SEPs that characterised depth, amplitude, and latency of negative components of the SEP using single electrodes [5,30,37]. Fluorescent staining of the recording site localised the maxima of the N1 component to lamina IV/V of the spinal dorsal horn, which is known to be innervated by Aβ-afferents encoding aspects of innocuous touch [61]. Following noxious electrical stimulation, evoked N2 and N3 components were observed at the same depth of the N1 component, with latencies consistent with Aδ-afferent and C-nociceptor conduction velocities [31]. These longer latency SEP components reflect the gradual recruitment of different afferents of the nociceptive system to higher stimulus intensities. Although nociceptive Aδ-afferents and C-nociceptors synapse onto nociceptor specific second order neurons in the superficial dorsal horn, these fibres and Aβ-afferents also synapse with WDR neurons in lamina IV/V, which suggests the encoding of innocuous and noxious stimuli is occurring at the site generating this spinal SEP [21,57,68,70]. Human studies of the N13 SEP have also proposed WDR neurons as one of the neural correlates of the response [44,54] because their central role as targets of descending modulatory systems that might explain how the N13 is modulated in experimental pain models [24,25,48,49].

Tapentadol exerts an analgesic effect via two distinct mechanisms: µ-opioid receptor (MOR) activation and noradrenaline transporter reuptake inhibition (NRI) [60]. In this study, tapentadol significantly reduced the amplitude of the N1 component following low-intensity stimulation and this was also seen with incremental Inn-Nox ramping electrical stimulation. The reduction in the amplitude of the N1 component suggests tapentadol is inhibiting Aβ-mediated spinal dorsal horn activity to stimulus intensities across the innocuous-noxious range. Tapentadol has previously been shown to inhibit wide dynamic range (WDR) neuronal firing evoked by brush and mechanical noxious stimuli, in both cancer pain and spared-nerve-injury rodent models [6,27]. However, these studies somewhat paradoxically reported that tapentadol did not modulate Aβ-mediated activity evoked by supramaximal electrical stimulation (Figure 1D of Falk et al 2015, and Figure 5 in Bee et al 2011).

Many WDR neurons are projection neurons which receive convergent inputs from multiple primary afferent classes [15,68,70]. In the present study, tapentadol significantly reduced the N1 SEP following Inn-Nox stimulation, however it did not affect the EC50 (the current required to evoke 50% of the peak amplitude). This suggests that tapentadol does not modulate the N1 SEP by reducing the excitability of Aβ-primary afferent neurons (which would be reflected in an increase of the EC50), but instead it acts via post-synaptic mechanisms. This could be via MOR mediated inhibition of WDR neuron or via NRI [6,27], enhancing endogenous analgesic effects from descending noradrenergic inputs from locus coeruleus [34,36]. Since pharmacological studies had shown that acute behavioural effects of tapentadol are inhibited by MOR antagonists and noradrenergic antagonists only counteract chronic behavioural effects, we suggest that N1 inhibition is via tapentadol action at postsynaptic MOR.

Voltage-gated sodium channels (VGSCs) determine the excitability and initiation of action potentials in primary afferent neurons and are a prime target for analgesic drug development [23,47]. This study has shown that lacosamide reduced N1 amplitude and increased EC50 in response to electrical stimulation, without attenuating the peak following stimulation in the noxious range. As a non-selective VGSC blocker, lacosamide reduces the excitability of all primary afferent neurons by prolonging the slow-inactivation state of VGSCs [11,58]. An increase in the threshold of peripheral neurons following dosing with lacosamide inhibits the generation of action potentials in response to low-intensity stimulation, as evidenced by a reduction in the N1 amplitude compared to baseline. This mechanism would explain how lacosamide increased the EC50 of the N1 SEP without altering its maximum amplitude. However, previous *in vivo* electrophysiological rat studies have shown that systemic lacosamide inhibited spinal dorsal horn activity at a latency consistent with c-nociceptor mediated spiking, but not in the Aβ-fibre mediated range [7]. The differences in stimulus intensity used between the studies could account for this, as Bee & Dickenson used cutaneous stimulation of the receptive field at an intensity three-times the c-nociceptor threshold. Such noxious electrical stimulation would have supramaximally activated Aβ-fibres, possibly masking any inhibitory effects of the drug. The present study observed an effect of lacosamide on the amplitude of the N1 SEP to low-intensity stimulation but not the Inn-Nox stimulation. Indeed, Bee & Dickenson reported that lacosamide reduces spinal dorsal horn neuronal firing to brush in a rat spared nerve ligation (SNL) model, indicating lacosamide is acting peripherally on Aβ-primary afferents, albeit only in SNL rats [7].These findings fit with how lacosamide is used to treat peripheral neuropathies [29,33].

Finally, pregabalin showed no effects on the amplitude of the N1 potential, despite it being previously reported to prevent the onset of capsaicin-induced modulation of the N13 in humans [44]. Spinal application of 100μg pregabalin in a rat model of neuropathic pain has been shown to have a significant antiallodynic effect to noxious-electrical stimulation and an associated inhibitory effect on the hypersensitivity of dorsal horn WDR neurons [20]. An explanation for this discrepancy may be found in the mechanisms by which pregabalin acts in the spinal cord to modulate neuronal excitability. It is thought that pregabalin acts through the α_2_δ subunit of voltage gated calcium channels (VGCCs) in the spinal dorsal horn, reducing the presynaptic release of glutamate [63]. Therefore, it is possible an upregulation of α_2_δ subunit of VGCC in the rat dorsal root ganglion of SNL rats results in a greater efficacy of pregabalin in the spinal dorsal horn [17,22]. Indeed, Di Lionardo et al observed no effect of pregabalin on the amplitude of the N13 SEP in the control site of a capsaicin-induced peripheral sensitisation model, however pregabalin did attenuate an increase from the sensitised site [44]. So, in models of sensitisation (acute or chronic), pregabalin interaction with the α_2_δ subunit of VGCC in the rat dorsal root ganglion is potentiated, thus increasing its effects on spinal dorsal horn neuronal activity, whereas in naïve models that phenomenon is not observed.

The innocuous stimulation paradigm was used to assess the pharmacokinetic (PK) relationship between drug exposure and the amplitude of the N1 SEP. In the tapentadol cohort, the amplitude of the N1 SEP in the 10mg/Kg group was significantly modulated after 10 minutes and plateaued at 40 minutes. Single i.p injection of 10mg/Kg tapentadol has been shown to be antinociceptive in rat pain models including the hot plate, tail flick, formalin test and diabetic polyneuropathy [65]. In the lacosamide cohort, the amplitude of the N1 was significantly reduced after 40 minutes and continued to reduce until the experimental endpoint. Lacosamide has previously been shown to exert analgesic effects in rodent models of muscular and neuropathic pain at doses of 10-30m/kg i.p [12,13]. Previous rat PK studies have shown that an hour after an intravenous dose of 30mg/Kg lacosamide, the blood plasma concentration is 120µM (∼30µg/ml) [36] which is in the same region seen with i.p dosing in the current study (135µM). There was no effect of pregabalin upon the SEP and the plasma concentrations of drug exceeded the EC50 value for pregabalin’s anti-neuropathic pain action in rat nerve injury models (9.3µg/ml to 16.2µg/ml=59-102µM) [9,10], indicating there was more than adequate exposure at the dose of 30mg/kg (145µM).

Our findings collectively establish a foundation for the N1 SEP to be used to assess novel analgesics by establishing it as a correlate of nociceptive processing in the dorsal horn. The next step will be to better define the functional neuronal populations activated in the spinal cord during the generation of the N1 SEP, which will further guide its application for testing novel analgesic compounds. Similarly exploring its modulation and utility as an assay in sensitised pain models would be important to support its translational role, particularly where aspects of human pain conditions can be best replicated.

## Acknowledgments

The authors wish to thank Dr Graeme Newton for manufacturing the custom bipolar Ag-AgCl stimulation electrodes. We would like to thank Mr Shrey Bijlani for his assistance with the histological processing and the Wolfson Bioimaging Facility for their assistance in image processing.

## Author contributions

Kenneth A J Steel, Methodology, Validation, Formal analysis, Investigation, Writing – Original Draft, Writing – Review & Editing, Visualisation, Project Administration. James P Dunham, Writing – Review & Editing, Supervision. Caterina Leone – Conceptualisation, Writing - Review & Editing. Andrea Truini – Conceptualisation, Writing - Review & Editing. Rolf-Detlef Treede, Conceptualisation, Writing - Review & Editing, Funding acquisition. Keith Geoffrey Phillps, Conceptualisation, Resources, Supervision, Funding acquisition. Jeffrey Krajewski, Supervision. Anthony Blockeel, Conceptualisation, Methodology, Supervision, Project administration, Funding acquisition. Anthony E Pickering, Methodology, Resources, Writing – Review & Editing, Supervision.

